# ‘Awake delta’ and theta-rhythmic hippocampal network modes during intermittent locomotor behaviors in the rat

**DOI:** 10.1101/866962

**Authors:** Nathan W. Schultheiss, Maximillian Schlecht, Maanasa Jayachandran, Deborah R. Brooks, Jennifer L. McGlothan, Tomás R. Guilarte, Timothy A. Allen

## Abstract

Delta-frequency network activity is commonly associated with sleep or behavioral disengagement accompanied by a dearth of cortical spiking, but delta in awake behaving animals is not well understood. We show that hippocampal (HC) synchronization in the delta frequency band (1-4 Hz) is related to animals’ locomotor behavior using a detailed analysis of simultaneous head- and body-tracking data. In contrast to running-speed modulation of the theta rhythm (6-10 Hz, a critical mechanism in navigation models), we observed that strong delta synchronization occurred when animals were stationary or moving slowly and while theta and fast gamma (55-120 Hz) were weak. We next combined time-frequency decomposition of the local field potential with hierarchical clustering algorithms to categorize momentary estimations of the power spectral density (PSD) into putative *modes* of HC activity. Delta and theta power measures from these modes were notably orthogonal, and theta and delta coherences between HC recording sites were monotonically related to theta-delta ratios across modes. Next, we focused on bouts of precisely-defined running and stationary behavior. Extraction of delta and theta power density estimates for each instance of these bout types confirmed the orthogonality between frequency bands seen across modes. We found that delta-band and theta-band coherence within HC, and in a small sample, between HC and medial prefrontal cortex (mPFC), mirrored delta and theta components of the PSD. Delta-band synchronization often developed rapidly when animals paused briefly between runs, as well as appearing throughout longer stationary bouts. Taken together, our findings suggest that delta-dominated network modes (and corresponding mPFC-HC couplings) represent functionally-distinct circuit dynamics that are temporally and behaviorally interspersed amongst theta-dominated modes during navigation. As such these modes of mPFC-HC circuit dynamics could play a fundamental role in coordinating encoding and retrieval mechanisms or decision-making processes at a timescale that segments event sequences within behavioral episodes.

## Introduction

Delta-frequency activity (1-4 Hz) is often observed in local field potential (LFP) recordings, but it is rarely analyzed in detail. This is in part due to the fact that during polymorphic slow-wave sleep (Steriade et al., 1993, Amzica & Steriade, 1998), delta waves correspond to periods of widespread cortical silence that encompass both principal cells and interneurons throughout cortical layers (Sirota & Buzsaki, 2005), giving the impression that neural computational processes are offline. Delta frequency activity is also evident in awake rodents (Bennett et al., 2013, Crochet & Petersen, 2006, Poulet & Petersen, 2008, Zagha et al. 2013), monkeys (Lakatos et al., 2005, Tan et al., 2014), and humans (Sachdev et al., 2015), but overt behavioral correlates of delta are lacking, and sensory stimuli or movement can elicit desynchronized states that supplant slow activity (Zagha et al. 2013, Tan et al., 2014).

During the slow oscillation of non-REM sleep, delta waves correspond to hyperpolarized down-states interspersed between HC ripple-associated and plastic phases of the cortical slow oscillation (Sirota & Buzsaki, 2005, Todorova & Zugaro, 2019). Interestingly delta activity has been shown to become increasingly prevalent during prolonged periods of wakefulness, even meriting comparison to neuroanatomically-local sleep in awake animals (Vyazovskiy et al., 2011), and delta waves are accentuated when sleep is finally possible after such deprivation (Vyazovskiy et al., 2009). Does this amplification of delta activity somehow ameliorate prolonged or excessive neural exertion, or is it otherwise related to the continuous accumulation of waking experience without the restorative punctuation that sleep provides? An important recent study found that, rather than absolute cortical silence during delta waves, a few neurons remain active, and neuronal ensembles active during delta waves bridge HC ripple-related replay events and subsequent plastic phases of the slow oscillation supporting memory encoding for recent experiences prior to sleep (Todorova & Zugaro, 2019). Perhaps then, delta is not exactly a sign of cortical sleep, but rather an indication that competing cortical ensembles need to be quieted in order to draw into greater relief the specific activities of neurons representing experiences to be encoded into memory.

In contrast to delta, the theta rhythm (6-10 Hz) in HC and connected structures is a well-organized oscillation that is critical for spatial navigation and episodic memory (Winson, 1978, Buzsaki, 2002, Buzsaki & Moser, 2013); shows frequency and power correlations to locomotor speed (Whishaw & Vanderworf, 1973, McFarland et al., 1975, Maurer et al., 2005, Hinman et al., 2016) and behaviors related to active attention and environmental sampling (Aghajan et al., 2017, Barth et al., 2018); exhibits functionally-relevant cross-frequency coupling to gamma oscillations (>25 Hz) (Canolty et al., 2006, Tort et al., 2009, Shirvalkar et al., 2010); and coordinates robust spiking within the HC (Okeefe & Recce, 1993, Foster & Wilson, 2007, Johnson & Redish, 2007), and its projection targets (Sirota et al., 2008, van der Meer & Redish, 2011), that is critical for learning, decision making, and representations of ongoing, remembered, or imagined future experiences (Richard et al., 2013, Pfeiffer & Foster, 2013, Redish, 2016, Schacter et al., 2007).

Running speed modulation of the HC theta rhythm is an integral mechanistic component of models of spatial navigation and episodic memory (Buzsaki & Moser, 2013, Schultheiss et al., 2015) and reflects, in part, the integration of sensory-motor information during locomotion (Bland & Oddie, 2001). Locomotion also modulates the firing rates and patterns of individual neurons in parahippocampal structures (Kropff et al., 2015, Hinman et al., 2016) as well as primary sensory cortices, and numerous recent studies have demonstrated that desynchronized and slow-oscillatory cortical states correspond to animals’ behavioral activity or quiescence (Neill & Stryker, 2010, Saleem et al., 2013, Reimer et al., 2014, Vinck et al., 2015, Zagha et al., 2013, Tan et al., 2014).

Analyses of oscillatory dynamics in HC during behavior have begun to reveal distinct operational states or ‘modes’ that are distinguishable in terms of the spectral content of the LFP and neuronal spiking (Barth et al., 2018, Kemere et al., 2013, Colgin, 2015, Amemiya & Redish, 2018, Zhang et al., 2019). Focusing on the delta and theta frequency bands, in the present study we asked to what extent modes of HC activity, defined by the spectral content of the HC LFP, reflect differences in network coupling and behavior. Using a combination of hierarchical clustering algorithms and time-frequency decomposition, we found that delta synchrony in HC was negatively related to speed, delta-dominated modes were orthogonal to theta modes in a manner mediated by locomotor activity, and both delta and theta modes were representative of intrahippocampal network coupling and mPFC-HC circuit states.

## Methods

### Subjects

Six adult male Long-Evans rats, weighing 600-700g at the time of surgery, were used in the study. Rats were individually housed in a climate-controlled vivarium and maintained on a reverse 12hr light/dark cycle. Daily experiments were conducted during the dark cycle active periods. Rats had free access to food and water with the exception that food was not available during a 1-2hr period prior to each session where rewards were given. All procedures were approved by the Institutional Animal Care and Use Committee at Florida International University.

### Implants

Rats underwent surgery to implant chronic recording electrodes. Four animals were implanted for HC recordings with 32-channel silicon probes arranged as tetrodes with 25 μm between adjacent electrode sites and impedances of 1.23 ± 0.32 MΩ. The eight tetrodes were distributed across four shanks at tip-to-tetrode depths of 78 μm and 228 μm (NeuroNexus A4X2-tet-5mm). Shanks were separated by 200 μm giving each probe a total length of 0.67 cm. During implantation, the long axis was oriented medial-lateral to sample proximal CA1 and CA2 of dorsal HC (Fig. 2A). Two animals were implanted for dual-site recordings with stainless steel wire electrodes (0.42 ± 0.25 MΩ) using custom designed (Autodesk Inventor) and 3-D printed implant bodies (3D Systems ProJet 1200) to house the electrodes coupled to an integrated electrode interface board. These implants comprised a grid of 18 electrodes targeting HC (2 × 2.5 mm, 4 electrode rows, 0.5 mm center-to-center) (Fig. 2B, top panel) and 14 electrodes arranged in two parallel rows (4 × 1 mm, 0.5 mm long-axis spacing) aligned to the rostrocaudal axis of mPFC (Fig. 2B, bottom panel).

### Surgical and post-operative procedures

General anesthesia was induced with isoflurane (5%) mixed with oxygen (0.8 L/min) and maintained throughout surgery at 1-4% as needed. Body temperature was monitored throughout surgery, and a Ringer’s solution (5% dextrose, 1 ml increments) was delivered at intervals of ~1 hour to maintain hydration. The hair was cut over the scalp and rats were placed in a stereotaxic device. Glycopyrulate (0.5 mg/kg, s.c.) was administered to assist respiratory stability, and ophthalmic ointment was applied to the eyes. Prior to incision, four injections of marcaine were made to the scalp (~0.1 ml, 7.5 mg/ml, s.c.). The pitch of the skull was leveled between bregma and lambda, two stainless steel ground screws placed in the left parietal bone, and four or five titanium support screws were anchored to the skull (given space constraints with dual-site and silicon probe implants, respectively). For silicon probe implants targeting dorsal HC, rectangular craniotomies were drilled to accommodate the probe shanks centered on coordinates AP −3.24 mm, ML 2.7 mm. For dual-site implants, craniotomies were shaped to accommodate the respective geometries of the electrode grids targeting HC (at AP −2.9 mm, ML 2.0 mm) and mPFC (AP 3.0 mm, ML 1.0 mm). After removal of the dura, implants were lowered on the stereotaxic arm until the electrode tips were just above the cortical surface. The ground wire was attached to the ground screws, and implants were lowered such that HC electrodes reached ~2.8 mm below the cortical surface and mPFC electrodes reached ~3.3 mm in depth. Sodium alginate was applied to the exposed surface of the brain within the craniotomy, and dental cement was applied to permanently affix the implant to the skull screws. Neosporin was applied to the skin surrounding the implant, and flunixin (2.5 mg/kg, s.c.) was administered for analgesia. Rats were removed from the stereotaxic device and allowed to wake up and rest on a heating pad until turgid, at which time they were returned to their home cages. Neosporin and flunixin were administered again the day following surgery, and rats were monitored closely for five days thereafter to ensure no negative symptoms emerged and that body weight was maintained. A 5-day regimen of daily Baytril injections (s.c.) was given during this time to preempt any post-surgical infection. At least one week was allowed for recovery from surgery prior to beginning experiments.

### Behavioral Apparatus

All experiments consisted of freely behaving rats in an open field environment (122×118×47 cm). The interior walls of the arena were lined with white vinyl; a white custom cutting board served as the floor; and all corners were sealed with white silicone. The arena was mounted on stilts 72 cm above the floor and the exterior was encased in grounded Faraday shielding. Two video cameras were used to track rats’ locomotion and other behaviors with a bird’s eye view. These were mounted near the ceiling, 28 cm apart from one another,150 cm above of the midline of the arena floor, yielding ~4.6 pixel/cm resolution in recorded videos and corresponding tracking data.

The entire behavioral apparatus was enshrouded with black curtains suspended from the ceiling and extending to the floor. Small gaps at the Velcro® closures at the corners between the four curtain panels were used to observe rats covertly during recording sessions to monitor that the head-stage, cables, and elastic tethers were not tangled, and to confirm that rats were awake during periods of relative inactivity. The white arena was illuminated dimly with red light from LED strips arranged in a square and mounted at the level of the video cameras. Outside the curtains, all lights were kept off during recording sessions save the experimenter’s computer monitor, which was not visible from the interior of the behavioral arena.

### Electrophysiological recordings

Throughout each experiment wide-band neural data were acquired with a Plexon OmniplexD recording system with A/D conversion at 40 kHz (16 bit). Briefly, a digital head stage processing subsystem (Intan, 32 channel) passed continuous wide-band data referenced against the ground screws (low cutoff = 0.7Hz) to be stored on a hard drive using PlexControl data acquisition software. For local field potential (LFP) analysis, the wideband data was down-sampled to 2kHz.

### Behavioral tracking

In parallel with neural recordings, Cineplex software (Plexon) was used to track and record (30 fps) the location of two clusters of LED florets (one green and one blue) that were offset ~2 cm laterally from attachments to the head-stage at ~3 cm above the rat’s head. Each LED cluster comprised three florets oriented upward and angled slightly outward. This construction reduced gaps in tracking data that resulted from a variety of behaviors including rearing and grooming in which rats oriented their heads away from the level plane or otherwise obscured the LEDs.

The physical and software settings for the two video cameras used for behavioral tracking were tuned independently to provide complementary data types for subsequent offline analyses. Whereas camera one was tuned to optimize tracking of the head-mounted LEDs with high color saturation and contrast against a near-black background; camera two was tuned to (1) maximize light sensitivity (sacrificing color saturation) allowing rats’ dark fur and body contours to be distinguishable against the white floor, while also (2) maintaining the ability to track the bright LED luminance peaks against the dimly illuminated environment. Raw videos from both cameras were recorded for all experiments, and online video tracking during each experiment yielded an output file for each camera containing the coordinates of the center of each LED cluster for each video frame. Video tracking frame captures were synchronized in real time to the simultaneously recorded neural data via a central timing board (Plexon).

After each experiment, the video file from camera two was retracked in Cineplex’s object contour tracking mode. Body location was calculated as the center of mass from difference images obtained by subtracting an image of the empty arena (captured just prior to placing the rat on the floor inside) from each frame of the video from camera two. Movement speeds were calculated from these body coordinates.

### Coregistration of LED and body tracking data

Following acquisition, all analyses of behavioral data were performed with custom-written Matlab scripts (MathWorks, Natick, MA). First, the coordinate system for camera one was mapped onto the coordinate system for camera two for each experiment, allowing LED and body tracking data to be integrated. These mappings were achieved as follows: First, we restricted the data to timepoints for which tracking data were complete (for both LEDs, both cameras, and body location) and free from identifiable tracking errors such as occasionally resulted from reflections of the LEDs on surfaces within the arena. Second, a 3^rd^ order polynomial function was fitted to each of the four sets of simultaneously-recorded 1-D coordinate pairs from the two cameras, defining the relationship, for example, between the x-coordinate of the blue LED on camera one to the x-coordinate of the blue LED on camera two. Using these functions, the precise LED tracking data from camera 1 was remapped onto the coordinate system for camera two allowing it to be integrated with the body tracking data.

We derived for each timestep the animal’s head direction (HD) relative to the environment (from the axis between the two head-mounted LED clusters); body direction (BD) relative to the environment (from the midpoint between the LEDs to the center of the body); head angle (HA) relative to the body (using the LED axis relative to BD); and the rotational velocity of the body.

### Definitions of locomotor and stationary bouts

To isolate bouts of locomotion from other behaviors including grooming and rearing, we first required that movement speed be maintained above 5 cm/sec for a minimum duration of 2.05 seconds with a minimum peak speed of 15 cm/sec. We then eliminated all instances where head angle exceeded 35 degrees or body rotational velocity exceeded 52.5 degrees per second at any time. Stationary bouts were required to meet the same minimum duration, HA, and rotational criteria and not exceed a maximum movement speed of 5 cm/sec.

### Rewarded behavioral sessions

To familiarize rats with the food rewards to be used, five Fruit Loops (TM), broken into individual rewards of ~1/4 loop, were given daily in rats’ home cages for five days prior to beginning these experiments. Subsequently, each rat in this study was run in two or more behavioral sessions lasting ~35 minutes each during which rewards were sporadically delivered. There was no explicit task in these behavioral sessions, but rewards were distributed sporadically one-at-a-time throughout the arena to motivate running to retrieve specific rewards as well as exploratory foraging generally. Rewards were distributed from above onto the arena floor using PVC pipes mounted at the ceiling outside the peripheral curtain enclosure and descending to terminate centrally adjacent to the video cameras. This approach was intended to minimize rats’ awareness of or attentiveness to the experimenter (which can be significant in similar behavioral paradigms using manual reward deliveries). None-the-less, some rats exhibited ‘anticipatory’ behaviors between reward deliveries, standing upright and fixating upward towards the reward delivery pipes.

Rats’ reward-seeking behaviors, including running speeds while retrieving rewards and the consistency and total duration of foraging during a session, as well as the frequency and duration of reward-irrelative behaviors, were variable across individuals and sessions, as well as within sessions. Thus, the temporal pattern and number of rewards delivered per session were not predetermined. Rather, the experimenter attempted to maintain two to five rewards distributed broadly across the OF at a given time while rats were actively foraging. Rewards were delivered infrequently (one to two per minute) while rats were relatively idle, allowing not more than ~10 to accumulate within the environment. No rewards were delivered while rats were actively grooming.

### Free exploration during non-rewarded behavioral sessions

Rats with silicon probe implants in HC (n=4) were also run for five non-rewarded sessions. These sessions were run prior to rewarded sessions while the environment remained novel and prior to developing expectations of reward in the arena context. During free exploration sessions, rats were placed on the arena floor and recorded without any intervention except for infrequent behavioral checks or to adjust cables or connections.

### Derivation of delta, theta, and gamma amplitude timeseries

To evaluate the correspondence of HC LFP dynamics in the delta-, theta-, and slow and fast gamma-frequency bands (1-4 Hz, 6-10 Hz, 25-55 Hz, and 65-120 Hz, respectively) to animals’ movement speeds, we first derived amplitude timeseries for each frequency band for each recording site, as follows: Peaks and troughs of each band-pass filtered LFP signal were identified; half-cycle amplitudes were calculated directly by subtracting each trough from the adjacent peaks; and these amplitude values, assigned to the median timepoints of the corresponding trough-peak intervals, were then resampled to 2 kHz to give timeseries paralleling the original LFP recordings.

### Movement speed modulation of HC rhythms

Timeseries for movement speed and the amplitudes of each frequency-band were mean-filtered with a two second sliding window using a one second step size. The two second window duration was chosen as a compromise between the relatively fast timescale of rats’ locomotor behaviors, e.g. bouts of running and interspersed pauses, and the reduction in number of oscillatory cycles (particularly for delta) contributing to amplitude measures for shorter time windows. We the regressed movement speed onto the four frequency bands of interest for each HC recording site from rewarded behavioral sessions chosen for each rat (n=164). We compiled across all regression models the distributions of beta values for each frequency band that was a significant predictor of movement speed (p<0.001). We also generated the distributions of pairwise correlation coefficients between each frequency band and movement speed, as well as between pairs of frequency bands.

### Spectral content and coherence of HC LFPs

The power spectral density (PSD) of HC recordings was estimated using Welch’s method for 2.05 second windows throughout each recording session. Raw PSDs were clustered hierarchically using MATLAB’s ‘linkage’ function into 16 groups based on frequency content below 25 Hz. This clustering approach was strongly influenced by PSD amplitude. We also clustered area-normalized versions of the same PSDs resulting in groupings emphasizing similarity of PSD waveforms independent of amplitude. In both cases, once cluster membership was determined, measures of delta and theta power were derived from the raw PSDs. Coherence estimates between each of six pairs of four simultaneously-recorded HC sites were obtained using MATLAB’s ‘mscohere’ function. The medians of these coherence estimates, derived for LFP timeseries for each instance of putative modes and behavioral bouts described above, were used for comparisons across conditions.

### Histology

After completion of all experiments, rats were anesthetized with isoflurane (5%) mixed with oxygen (800 ml/min) and marking lesions were made with a NanoZ (Plexon) to deliver 10μA for 10s at each of the 32 electrode locations. About an hour later, rats were transcardially perfused with 100 ml phosphate-buffered saline (PBS), followed by 200 ml of 4% paraformaldehyde (PFA, pH 7.4; Sigma-Aldrich, St. Louis, MO). Brains were post-fixed overnight in 4% PFA and then placed in a 30% sucrose solution for cryoprotection. Frozen brains were cut on a Leica CM3050 S cryostat (40 μm; coronal) and Nissl-stained. Marking lesions were mapped onto plates of the Paxinos & Watson Atlas (2018).

## Results

Focusing on delta and theta frequency bands, we investigated the spectral content of HC LFPs while rats freely navigated a large ‘open field’ arena (122 cm × 118 cm) in both non-rewarded and sporadically-rewarded (dropped Fruit Loops®) session types (Fig. 1). Locomotor coverage of the arena during representative sessions of each type is depicted in Fig. 1A&B, illustrating that the availability of rewards increased locomotor activity.

**Figure 1.**
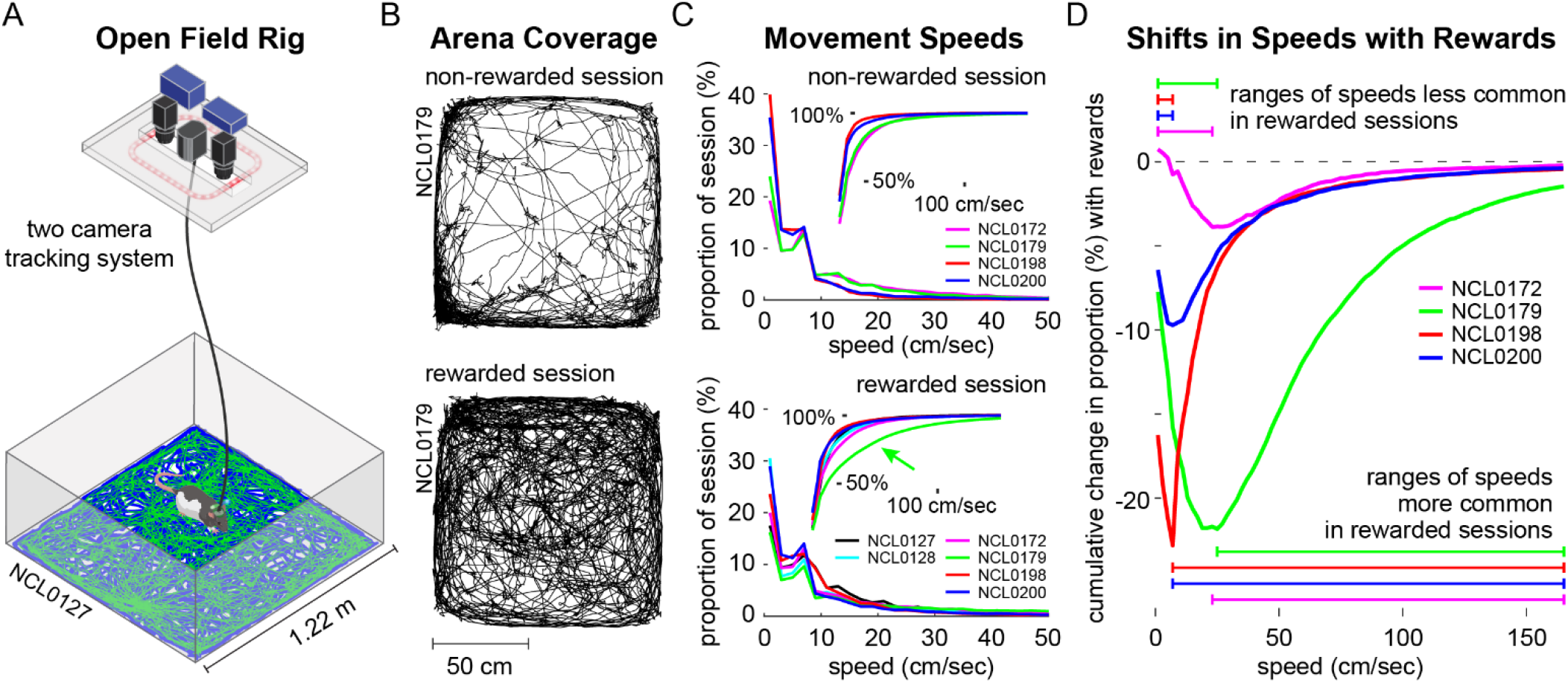
Locomotor behavior during free exploration and foraging. **A.** Open field behavioral arena with separate cameras for tracking body contour and head-fixed LEDs. **B.** Examples of locomotor paths taken by the same rat during a non-rewarded session (**top**) and a rewarded session (**bottom**). **C.** Speed occupancy during non-rewarded (top) and rewarded sessions (bottom) for each rat. Insets show speed occupancy as cumulative proportions of session time across increasing movement speeds, e.g. rat NCL0179 spent a greater proportion of the rewarded session at faster movement speeds than did the other rats (green arrow). **D.** Changes in speed occupancy between non-rewarded and rewarded sessions, shown as the difference in proportion accumulated across speeds. Negative peaks in these curves reflect the speed below which rats spent less time during rewarded sessions in favor of faster speeds.

**Figure 2.**
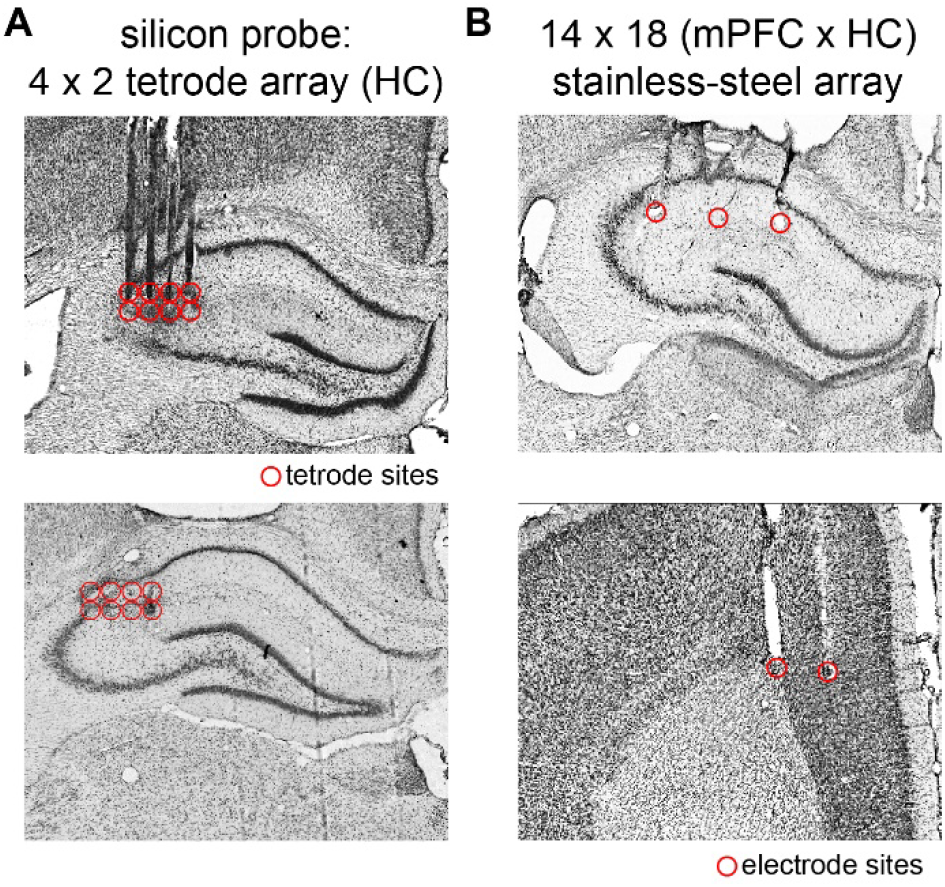
Local field potential recording sites in HC and mPFC. **A.** Representative brain slices through HC (**top** and **bottom**) showing tracks left by silicon probe shanks and recording sites (red circles) near the junction of proximal CA1 and CA2. **B.** Representative brain slices through HC (**top**) and mPFC (**bottom**) showing stainless steel wire electrode tracks and recording sites.

Overall behavior. During both non-rewarded and rewarded sessions, rats were more often stationary (~20-40% and ~15-30% of sessions, respectively) than walking or running at any other particular speed (Fig. 1C). As has been previously observed, a consistent peak in speed occupancy was evident at ~7 cm/s (e.g., Wirtshafter & Wilson, 2019) reflecting the prevalence of slower exploratory behaviors. Across all sessions the average distance traveled was 36.46 ± 14.45 m, with more distance covered in rewarded (40.19 ± 15.55 m) compared to non-rewarded sessions (29.38 ± 8.67m; t_(52.8)_ = −3.31; P = 1.7×10^−3^). The corresponding average body movement speed across all sessions was 12.9 ± 4.8 cm/s, with faster movements during rewarded (14.1 ± 5.3 cm/s) compared to non-rewarded sessions (10.64 ± 2.5 cm/s; t_(52.7)_ = 3.26; P = 1.9×10^−3^). A repeated measures comparison of average speed during consecutive non-rewarded (10.8 ± 3.1 cm/s) and rewarded session pairs (17.0 ± 8.5 cm/s) (n=4 rats) also indicated faster body movements (and reduced time spent stationary) during rewarded than non-rewarded sessions (t_(7)_ = 2.7, P = 3.0×10^−2^) (Fig. 1D), and movement speeds were further elevated during the second rewarded session (17.2 ± 7.1 cm/s) compared to the first (14.6 ± 7.4 cm/s; t_(5)_ = 2.6, P = 4.6×10^−2^).

As is typical in open field experiments, rats tended to stay close to walls and corners of the arena (i.e., thigmotaxis, as in Fig. 1B, top) with an average across all sessions of 76.5 ± 20.6% time spent in the areas nearer to a wall and 23.5 ± 20.6% time in an equivalent central area. In rewarded sessions (the first of which was subsequent to the last non-rewarded session for each of four rats) more time was spent in the central space (29.8 ± 21.5%) compared to non-rewarded sessions (11.6 ± 12.17%; t_(52.7)_ = 4.01; P = 1.9×10^−4^), reflecting routes taken for retrieval of rewards and diminished anxiety or environmental novelty during the later sessions.

### Incidence of precisely-defined behavioral bouts

In addition to spatial occupancy and movement speeds measured from body tracking data, we derived animals’ body orientation, head angle, and rotational velocity by integrating tracking data for head-mounted LEDs with body tracking data (Fig. 1A). We found that rats’ head angle exceeded 35 degrees (as when looking to the side or turning) during 13.76 ± 5.8% of each session, and rats’ rotational velocity exceeded 52.5 degrees/sec during 15.10 ± 5.6% of each session. These proportions did not differ between session types (p = 0.59 and 0.69, respectively). Using these thresholds in conjunction with movement speed and duration criteria, we were able to precisely identify bouts of locomotor and stationary behavior, while excluding the majority of other fast movements (e.g. grooming) or movements ‘*in place’* (e.g. rearing). The number of locomotor bouts exhibited per session varied widely between rats as well as across sessions (>2 sec duration: 5 to 45 bouts, mean = 15.5; >1 sec duration: 24 to 239 bouts, mean = 94.5), but neither the incidence nor the maximum duration of locomotor bouts (4.1 ± 0.76 and 4.3 ± 1.0 sec for rewarded and non-rewarded sessions, respectively) differed between session types using pairs of repeated measures (n=4 rats; P_(2sec)_=0.53 and P_(1sec)_=0.27; P_(max)_=0.82). However, average movement speeds during locomotor bouts were modestly faster in rewarded sessions (29.2 ± 7.9 cm/sec) than nonrewarded sessions (25.6 ± 5.9 cm/sec; tstat_(7)_=2.57; P=3.7×10^−2^; tstat_(7)_ = 2.77; P=2.7×10^−2^, respectively). Taken together, our behavioral analyses suggest idiosyncrasies in rats’ propensity to explore, but consistency in the temporal features of locomotor bouts and relationships to reward.

### Recording sites in HC and mPFC

HC electrodes were targeted to the proximal CA1-CA2 juncture (Fig. 2A & 2B, top panel) of the right dorsal HC (A/P: −2.52-3.72 mm, M/L: 1.00-3.00 mm). The majority of HC marking lesions were found in stratum lacunosum moleculare or stratum radiatum (M/L: 1.00-3.00mm). Recordings from electrodes in the corpus callosum or CA3 were not analyzed. Marking lesions at recording sites distributed along the rostrocaudal extent of mPFC (Fig. 2B, bottom panel) were found primarily in the deep layers of anterior cingulate cortex and dorsal prelimbic cortex (A/P: 4.68-2.16mm, M/L: 0.8-1.4mm). In one rat, the lateral row of mPFC electrodes invaded motor cortex and was excluded from further analysis.

### Spectral content of HC LFPs during navigation

The power and peak frequency of the HC theta rhythm (6-10 Hz), as well as theta-gamma cross-frequency coupling, have been shown to covary with an animal’s running speed in several behavioral paradigms (Maurer et al., 2005, Hinman et al., 2016, Richard et al., 2013, Sheremet et al., 2019), and the theta rhythm is an important mechanistic component of episodic memory and spatial navigation models (Schultheiss et al., 2014). As expected, our HC LFP recordings consistently showed a prominent theta rhythm as rats navigated the open field (Fig. 3A-C), and theta power was markedly elevated during faster locomotion (Fig. 3D-I). Spectrograms shown in Figure 3 (D, F, & H) illustrate the time-varying spectral content of HC LFPs and highlight the correspondence of theta power (warmer colors) to faster movements (white traces). Moreover, theta harmonic power (~16 Hz) often appeared in spectrograms coincident with peaks in movement speed (open stars). In striking contrast, significant power in the delta frequency band (1-4 Hz) often developed immediately when animals became still after periods of movement (filled stars in Fig. 3; Supp. Fig. 1). These delta events appeared in the spectrogram as tails descending from the theta band in tandem with pauses in rats’ locomotor activity (white traces). Although elevated delta power typically persisted while animals remained stationary, LFP voltage traces generally did not show a stable, rhythmic delta oscillation with continuous phase progression across successive cycles. See Supp. Fig. 1A and Fig. 3 A&B showing slow voltage fluctuations without well-organized periodicity.

**Figure 3.**
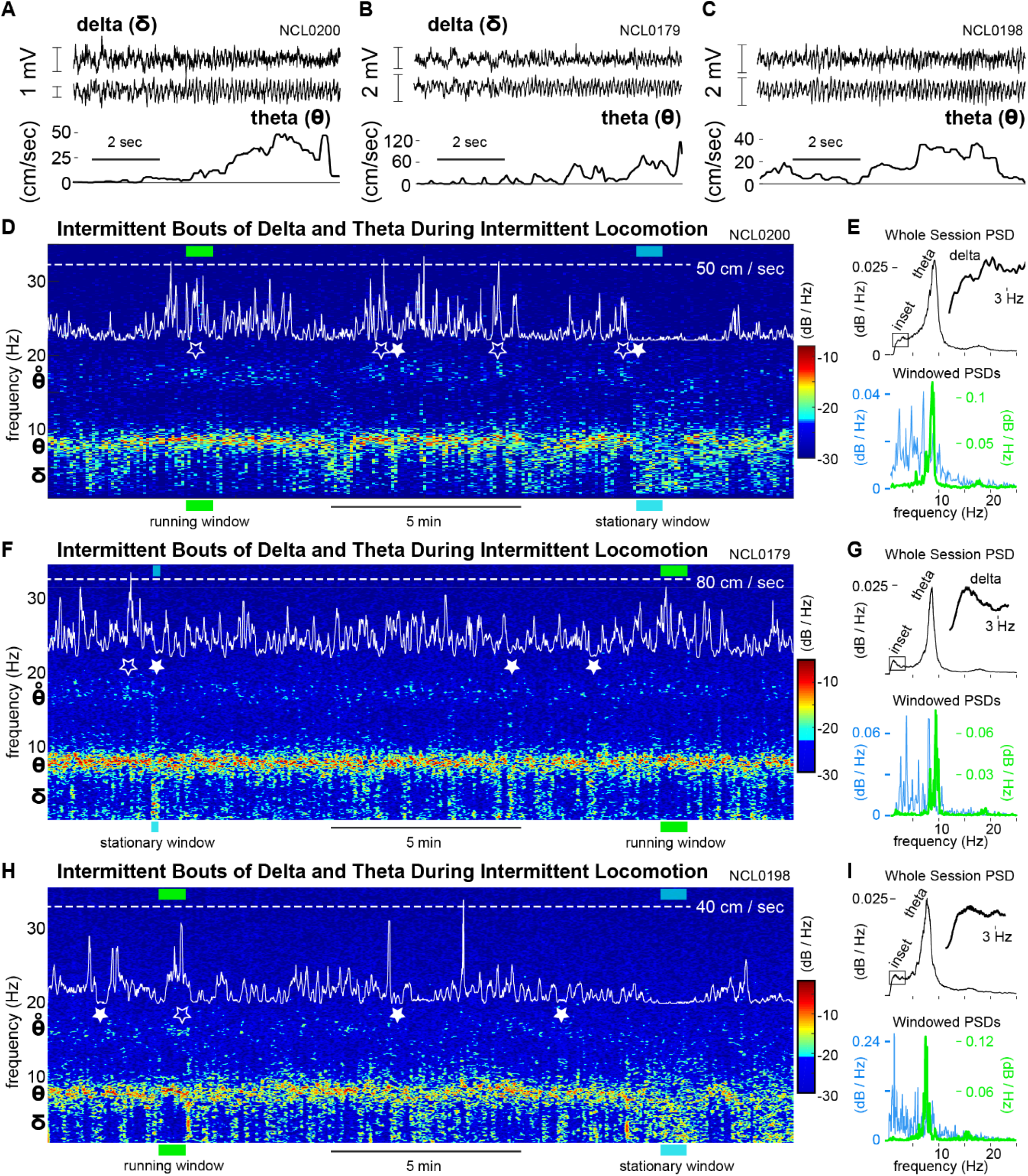
Spectral content of HC LFPs during navigation. **A-C.** Pairs of representative, simultaneously recorded HC LFPs (top two traces) and corresponding movement speeds (bottom trace) for three rats. **D.** Spectrogram of the HC LFP showing the theta rhythm (6-10 Hz) prominently throughout the session (warm colors). Movement speed is overlaid on the spectrogram (white trace) to allow comparison of running speed peaks (e.g. open stars, accompanied by theta-harmonic power) and stationary behavior (filled stars) to the spectral content of the LFP. **E.** Power spectral densities for the whole recording session (top) and for representative windows (36 seconds) of running and stationary behavior. Timing of these windows within the recording session is indicated with green and cyan bars in **D**. **F-I.** As **D**&**E**, for two additional rats.

Delta-band synchronization typically appeared in the power spectral density (PSD) for a given session as either a small distinct peak or as an asymmetric shoulder to the theta peak (Fig. 3 E, G, & I, top panels). These PSDs are equivalent to integrating the entire spectrogram over time into a single spectrum. In contrast, PSDs for shorter epochs (≤36 seconds) often showed either a sharply-tuned theta peak and very little delta power when animals were running fast, or elevated delta power without a prominent theta peak when animals were stationary (Fig. 3 E, G, & I, bottom panels). This observation led us to ask whether delta-band synchronization is (1) related to locomotor behavior, and (2) coupled to rhythmic activity in other frequency bands.

To evaluate the hypothesis that HC delta synchrony (in addition to the theta and fast gamma rhythms) is modulated by movement speed, we derived for each LFP channel an instantaneous-amplitude timeseries for each frequency band (Fig. 4A). The derivation of these timeseries used interpolation between the times of peak-to-trough measurements for each cycle at each frequency, yielding smoothly varying amplitude estimates with a constant sampling timestep. We then calculated the pairwise correlation coefficients between movement speed and the amplitude of each band (Fig. 4B), and we asked whether the distributions of r-values (across all recording sites in all rats) were different than zero. Conceptualized in this way, we found that delta, theta, and fast gamma were all related to movement speed (Fig. 4C). Theta and fast gamma amplitudes both covaried positively with movement speed, but strikingly, delta showed the strongest relationship to movement speed (p=8.97×10^−65^) with almost no incidence of non-negative r-values across all HC recording sites (<3%, top left). Thus, whereas the theta and high gamma rhythms were strongest during fast running, delta power was strongest when animals were stationary or moving slowly and was diminished or absent during running. Moreover, using a four-predictor regression of movement speed onto delta, theta, and slow and fast gamma amplitudes, we found a significantly negative relationship of delta amplitude to speed and significantly positive relationships of theta and fast gamma to speed (Fig. 4D). Interestingly, the amplitude of slow gamma in our recordings showed conflicting relationships to movement speed in the two correlation analyses, and many fewer recordings sites exhibited speed modulation of slow gamma than of the other rhythms in the multiple regression.

**Figure 4.**
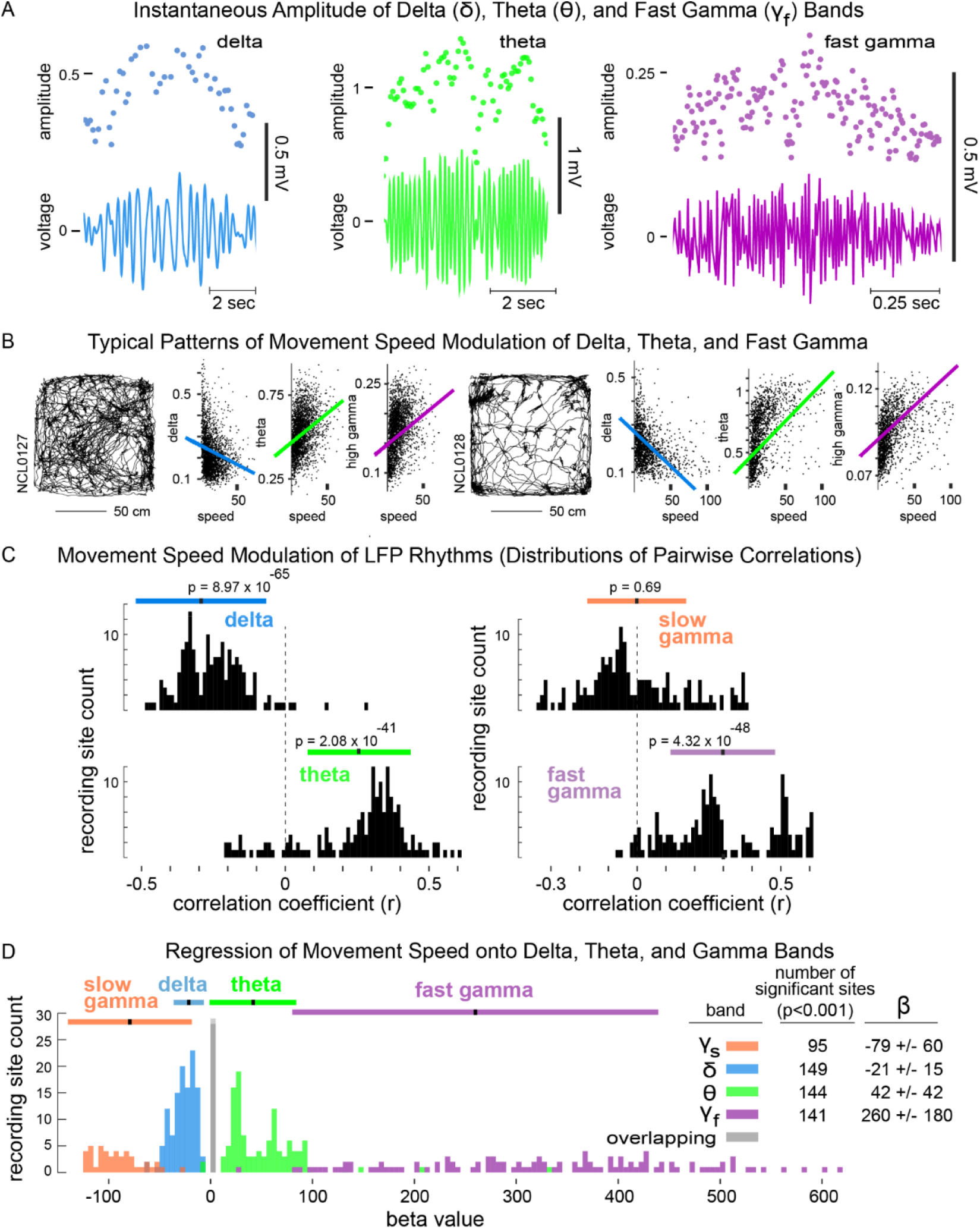
Movement speed modulation of delta, theta, and gamma amplitudes. **A.** Instantaneous amplitudes of delta, theta, and gamma were derived from peak-to-trough measurements (dots above) of corresponding band-pass filtered LFPs (voltage traces below). **B.** Scatterplots of instantaneous amplitudes of delta, theta, and high gamma, and overlaid best-linear-fits, corresponding to rats’ locomotor trajectories (left) during two representative sessions. **C.** Distributions across all rats and recording sites of pairwise correlation coefficients relating the instantaneous amplitude of each frequency band to movement speed. **D.** Distributions of β-values for significant predictors (p’s<0.001) of movement speed from regressions onto instantaneous amplitudes of delta, theta, and slow and fast gamma bands. The inset table shows the number of recording sites for which each band was a significant predictor and the corresponding mean β-values and standard deviations.

Next, to assess amplitude-amplitude cross-frequency coupling between rhythms, we inspected the distributions across recording sites of correlation coefficients between each pair of frequency bands (Fig. 5). We found that (1) overall, delta and theta amplitudes exhibited a weak tendency toward negative covariation based on the distribution of r-values; (2) delta and fast gamma showed a bimodal distribution of r-values consisting of a prominent negative peak and a secondary peak centered near zero (indicated with arrowheads); and, (3) theta and fast gamma also showed a bimodal distribution of r-values, but both peaks occurred at distinctly positive values suggesting the possibility that different theta-fast gamma coupling modes (strengths) were expressed at different subsets of recording sites. To unpack the composition of these r-value distributions, we also inspected the relationships between frequency bands for each recording site independently. Scatter plots of amplitude measurements for each pair of bands revealed a diversity of complex relationships (Fig. 5, all insets). Representative examples from two rats shown in Figure 5 illustrate that delta and theta amplitudes could be either (1) orthogonal, such that larger values and the preponderance of variance in each band occurred at the smallest amplitudes of the other (A, top inset), or (2) weakly anticorrelated (A, bottom inset).

**Figure 5.**
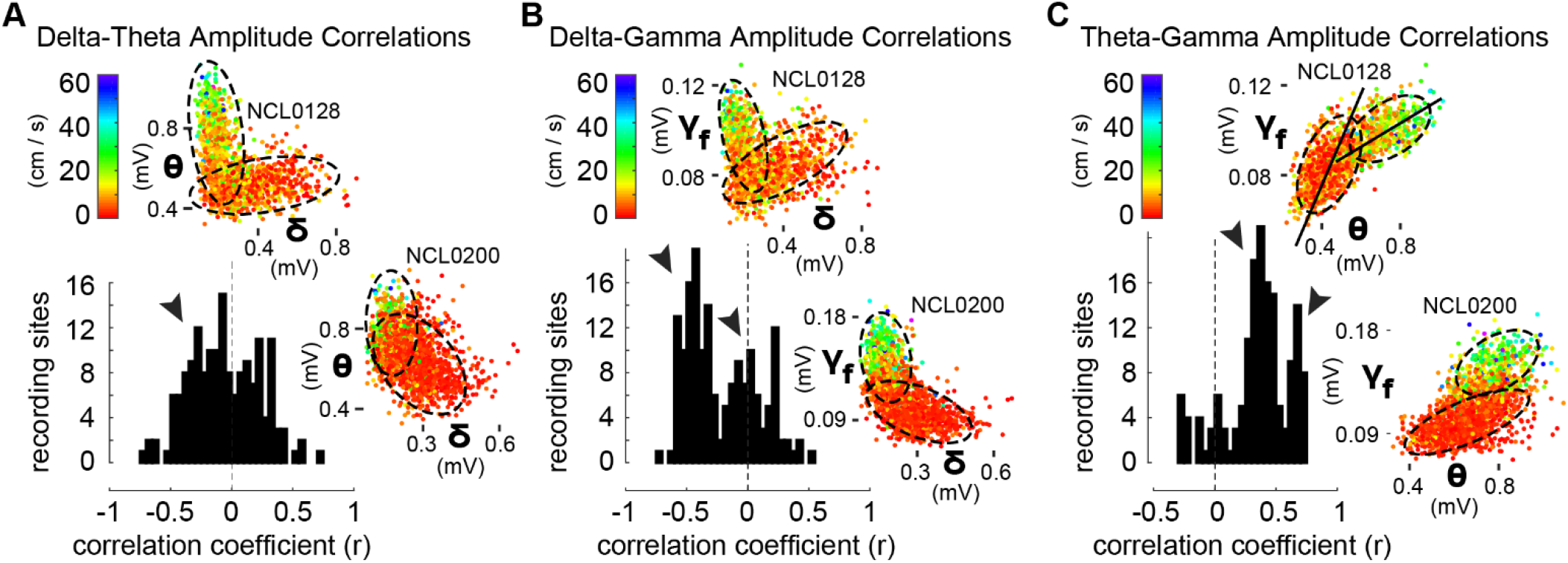
Multimodal distributions of cross-frequency amplitude couplings comprise diverse putative modes. **A.** Distribution across HC recording sites of r-values for pairwise correlations between instantaneous amplitudes of delta and theta (black histogram). Scatterplots inset above and at right illustrate for typical orthogonality and anticorrelation between bands, respectively, for one recording site from each of two rats. The weakly-negative histogram peak (arrowhead), merges all individual cases as in the insets. Visually-distinguished delta- and theta-dominated putative modes (dashed ellipses) corresponded to behavioral state (color code for movement speed at top left). **B.** Bimodal distribution of r-values between delta and fast gamma amplitudes (black histogram, arrowheads indicate one prominent negative peak and one peak at zero), and representative scatterplots of delta and fast gamma amplitudes for the same recordings as in insets in **A**. Orthogonality, and within-mode anticorrelation during running (red mode for rat NCL0200) are evident in visually-defined putative modes (ellipses). **C**. Bimodal distribution of r-values between theta and fast gamma amplitudes (black histogram, arrowheads indicate two prominent positive peaks), and representative scatter plots for the same recordings as in insets in **A**&**B**. Insets show globally-positive correlations between theta and fast gamma amplitudes, but these comprise visually-defined putative modes (ellipses) characterized by specific slopes between bands (long axis of ellipses, as in top inset).

Coalescence of these different delta-theta relationships is consistent with the modest negative peak in the r-value distribution. Scatterplots of delta and high gamma amplitudes showed similar relationships (B, insets), whereas theta and high gamma were positively correlated in most cases (C, insets). Importantly, these plots illustrate that correlated cross-frequency relationships are heterogenous across rats and recording sites, and each may comprise one or more distinct functional modes of cross-frequency coupling (suggested by dashed ellipses). Stationary behavior and running (color code for movement speed in all insets) consistently corresponded to different regions within 2-D cross-frequency amplitude spaces (insets), and amplitude relationships between bands exhibited characteristic slopes for each putative mode (i.e. the slope of the major axis of each ellipse, as in C, top inset).

Taken together, our analyses of speed modulation of HC LFP bands implicate the delta frequency band as a potentially important complement to theta and fast gamma rhythms in neural algorithms for navigation behavior. Significant further investigation is needed, however, to identify and characterize possible contributions of HC delta synchrony to (1) spatial coding during navigation, (2) retrieval of remembered information to guide navigation of previously experienced contexts, or (3) encoding of ongoing spatial and behavioral trajectories into episodic memory. A key first step is to clarify whether instances of strong delta synchrony and instances of strong theta synchrony represent distinct functional modes of network activity that are mutually-exclusive, or whether the two frequency bands can vary smoothly in opposition to one another, one waxing as the other wanes.

Given considerable behavioral variability between rats in our experiments and the dependence of delta- and theta-frequency synchrony on navigation behaviors, we devised a simple, behaviorally-robust analytic strategy to categorize putative modes of HC network activity based on the power spectral density (PSD) of the LFP (Fig. 6A-C). Briefly, we applied hierarchical clustering routines to the power spectra (0.7-25 Hz) for two second windows of the HC LFP throughout each behavioral session (i.e., the spectrogram). This general strategy for unsupervised categorization of network modes can be implemented in numerous ways using different clustering algorithms and parameter settings to focus on different aspects of the relationships between animals’ behavior and LFP spectra. We chose parameters to balance the frequency resolution of PSDs with the ability to analyze tracking data at behavioral timescales.

**Figure 6.**
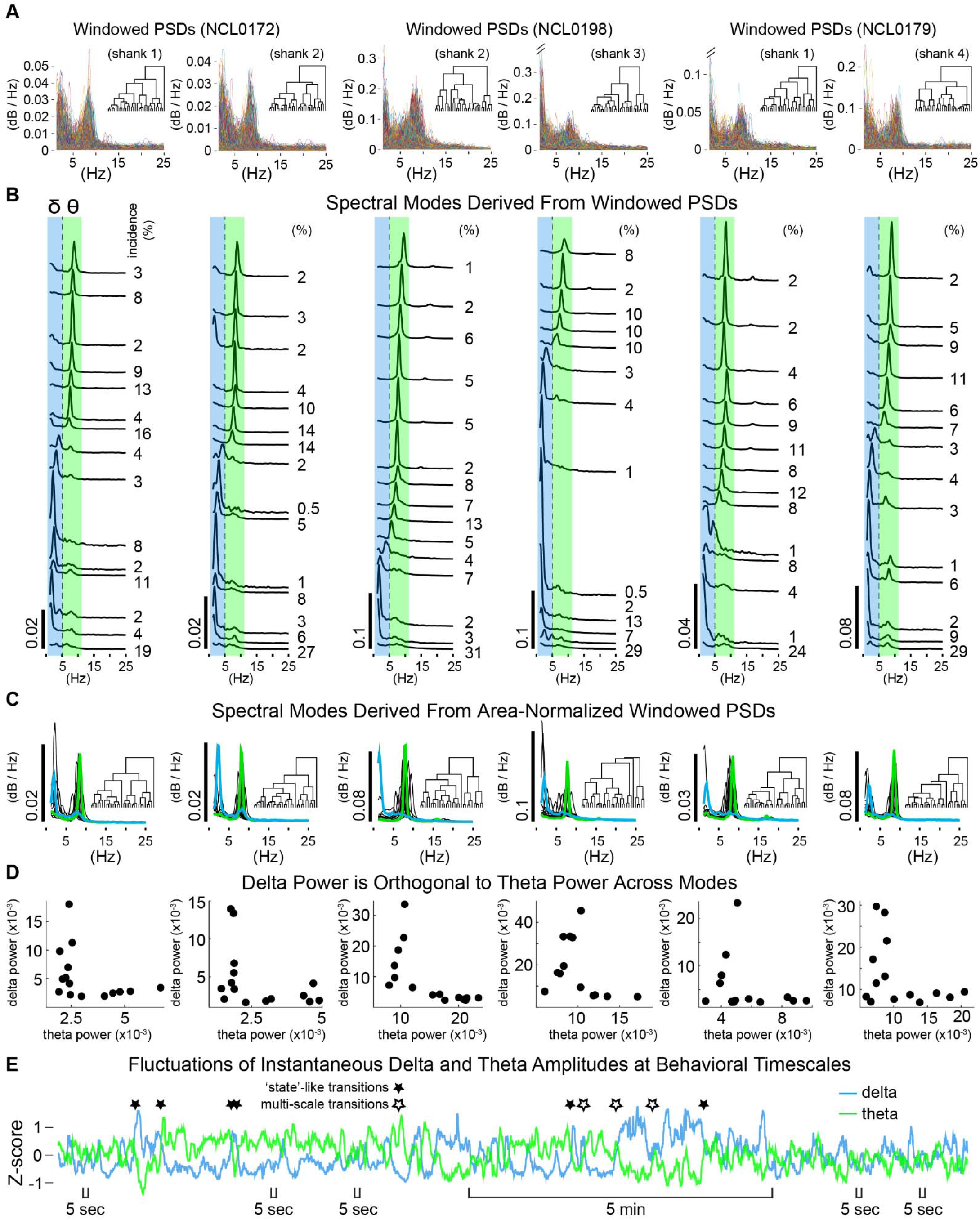
Spectral modes of HC network activity from hierarchical clustering of PSDs. **A.** Stacks of all power spectral densities (overlaid) from two second windows of HC LFPs. Examples are shown for two simultaneously recorded channels from each of three rats. Inset dendrograms show the cluster hierarchies. **B.** Spectral modes defined as the average PSD from each cluster of windowed PSDs in **A**. Larger amplitude modes were primarily evident central to the theta frequency band (green) or within the delta band (blue). The incidence of each mode (% of session) is indicated to the right. **C**. Spectral modes defined conservatively using area-normalized windowed PSDs (dendrograms inset). Representative delta-dominated and theta-dominated modes (average PSDs for example clusters) shown in blue and green, respectively. **D**. Measures of average power within delta and theta frequency bands were orthogonal across modes. Each dot represents delta and theta measures for one mode. **E**. Example time-course of delta and theta showing near-simultaneous fluctuations of the two frequency bands in opposite directions. Several sharp, opposite fluctuations (e.g. filled stars) are consistent with state transitions, but note occasional instances where delta and theta covary (positively) at a fast timescale while diverging over a slower timescale (multi-scale transitions, open stars). Also note that opposite fluctuations in delta and theta can be large or small and take place rapidly. (Five second scale bars indicate a few of many instances where smaller, fast fluctuations in delta and theta directly oppose one another.)

Figure 6A shows ~4000 superimposed time-windowed PSDs (for timepoints throughout behavior sessions) for two representative LFP channels from each of three rats. Each spectral mode derived by clustering windowed PSDs in this manner represents a vector of logical values (0s and 1s) indicating whether, at a given moment in time, the LFP showed a particular pattern of spectral content. That is, applying clustering routines to the spectrogram returned the times at which each spectral mode occurred. Using these logical values then allowed evaluation of whether specific aspects of neural activity (e.g. power in a specific band) or of behavior (e.g. running speed) varied between those sets of moments.

We found that delta power and theta power were strikingly orthogonal across spectral modes, which we defined as the average PSD for each cluster (Fig. 6B-D). This orthogonal relationship was consistently pronounced across rats and recording sites, amplifying our previous demonstration that larger amplitudes of either delta or theta typically occurred at times when synchrony in the other band was weak. Figure 6B shows the spectral modes derived from the windowed PSD ‘stacks’ in Figure 6A, sorted loosely according to peak frequency. Spectral modes almost always consisted of a single primary peak occurring at a frequency within the delta band or within the theta band, but not between delta and theta, particularly if the peak was of substantial amplitude (Fig. 6B). We also applied our clustering routines to stacks of area-normalized (<25 Hz) windowed PSDs. This manner of classifying spectral modes emphasized the relative power between frequency bands allowing the waveform of lower amplitude PSDs to contribute to mode classifications. With this more-conservative approach, we found a similar selectivity of spectral modes for either delta or theta (Fig. 6C). Note the consistent absence of modes with peaks between the delta and theta bands, as well as the absence or diminutiveness of secondary peaks for individual modes (Fig. 6C, green or blue traces). Figure 6D illustrates delta-theta orthogonality as the ‘elbow’ in plots of the relative power for the two bands across modes.

Delta and theta amplitude timeseries (as in Fig. 6E) showed that transitions between delta and theta often occurred rapidly, within a few seconds (as in Supp. Fig. 1), indicating that delta and theta modes could alternate during behavior sufficiently fast to implement different aspects of spatial encoding, decision, or memory processes. Fast ‘state-like’ transitions, whereby synchrony in one frequency band was replaced suddenly by synchrony in the other band were common (filled stars); but interestingly, these transitions were not always clean. In some instances, delta and theta fluctuated together (correlated positively) at a fast timescale, despite ongoing divergence between the two at longer timescales. Such ‘multi-scale transitions’ could be spurious or may represent nested neurophysiological phenomena of particular consequence. It will be important to determine what the impact of brief bouts or prolonged epochs of correlated delta and theta would be for encoding and plasticity in HC networks.

To begin to ask whether spectral modes are functionally distinct, we first evaluated whether animals’ behavior differed between times at which different modes occurred. We found that average delta power across spectral modes (classified using either unnormalized or normalized PSDs) was orthogonal to animals’ movement speeds (Fig. 7A&B, left panels). Theta power across spectral modes was strikingly correlated to movement speed, however, and this relationship was consistent across rats and recordings sites (t_(23)_=5.16; P=3.14×10^−5^, Fig. 7A&B, right panels). Next, we evaluated intra-HC phase coherence (between pairs of recording sites) as a function of spectral modes. Figure 7C shows the coherence spectra for each mode from a representative session. Note that coherence spectra with high delta-band coherence exhibited the least theta-band coherence (blue traces), and spectra showing high theta coherence had weak delta coherence (green traces). Lastly, using the ratio of theta power to delta power (TDR), a common neuropsychiatric diagnostic tool, we found that intrahippocampal delta coherence across modes consistently declined with increasing TDR (t_(23)_=−5.97; P=4.35×10^−6^), whereas theta coherence increased with increasing TDR (t_(23)_=4.912; P=5.8×10^−5^, Fig. 7D, Supp. Fig. 2).

**Figure 7.**
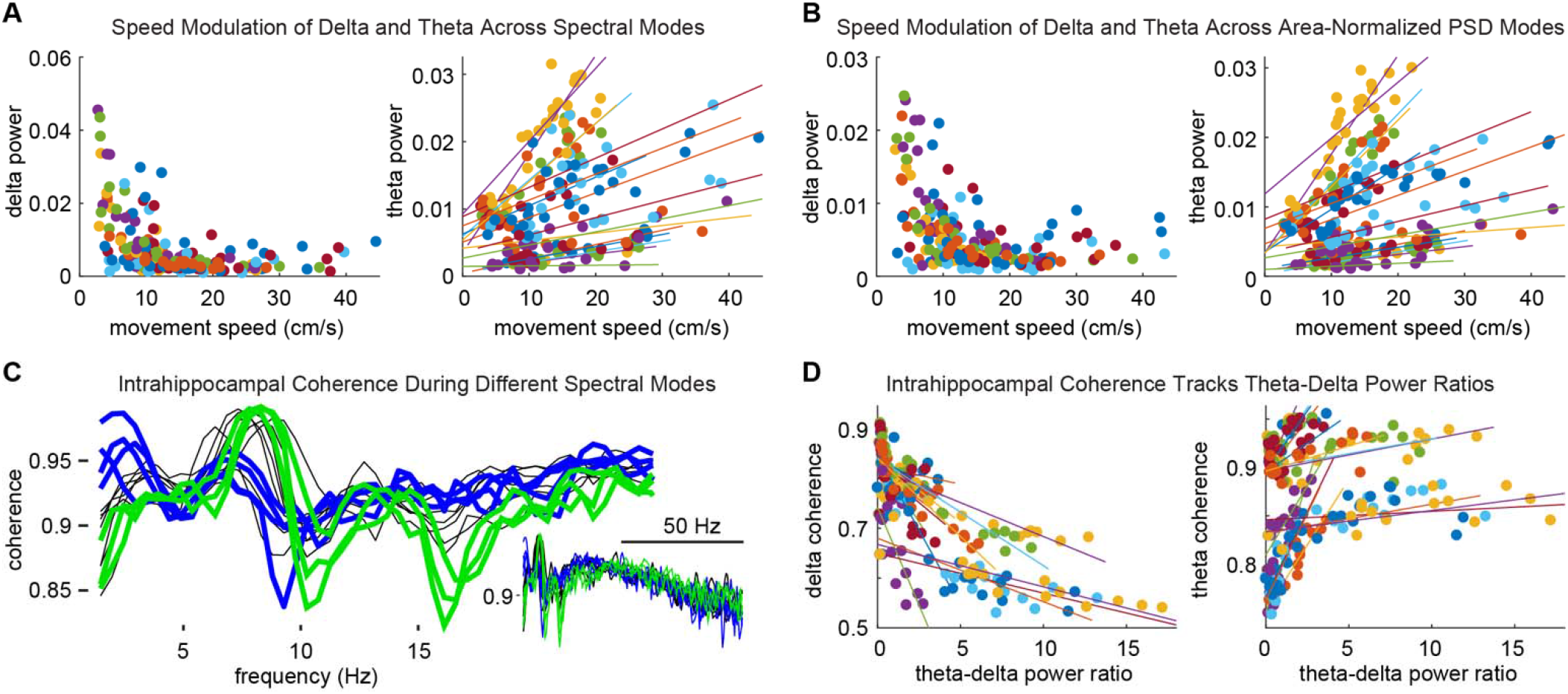
Spectral modes track behavior and network state. **A.** Modes with elevated delta power were evident when animals were stationary or moving slowly, whereas modes corresponding to running had minimal delta power (left). In contrast, theta power consistently scaled with running speed across modes. The different color represents different electrode channels from different rats (in A, B & D). **B**. Delta and theta power across modes corresponded to animals’ movement speeds as in **A**, using area-normalized mode definitions. **C**. Intrahippocampal coherence spectra for modes from a representative session. Blue traces highlight modes with high delta-frequency coherence and diminished theta coherence. Modes exhibiting high theta coherence between HC recording sites had diminished delta coherence. **D**. Across modes, delta coherence decreased monotonically (left) and theta coherence increased to ceiling (right) with increasing theta-delta ratios.

We have shown so far that across an entire recording session the amplitude of delta, theta, and gamma frequency bands are all related to movement speed, and band-pass amplitudes are coupled in a speed dependent manner. We have also showed that classification of HC network modes, purely by clustering power spectra, yields groupings that correlate with movement speed and intrahippocampal coherence, indicating that these modes are functionally distinguishable. In a last series of analyses, we refined our analysis of behavioral tracking data to isolate bouts of locomotor and stationary behaviors (Fig. 8) and exclude grooming, rearing, and other behaviors that incorporate fast rotational velocities or angling the head to either side (Fig. 8A). Figure 8 illustrates our analysis of tracking data used to identify running and stationary bouts. As expected, we found that delta power in HC was elevated when rats paused between runs even briefly and remained elevated while rats remained stationary. Theta power was elevated during running bouts and closely tracked average and peak speeds across running bouts. For the two rats implanted for dual site recordings from HC and mPFC (mPFC), we found that delta coherence between mPFC and HC was elevated during stationary bouts (p=1.6×10^−4^) and theta coherence was elevated during running (p=9.8×10^−27^).

**Figure 8.**
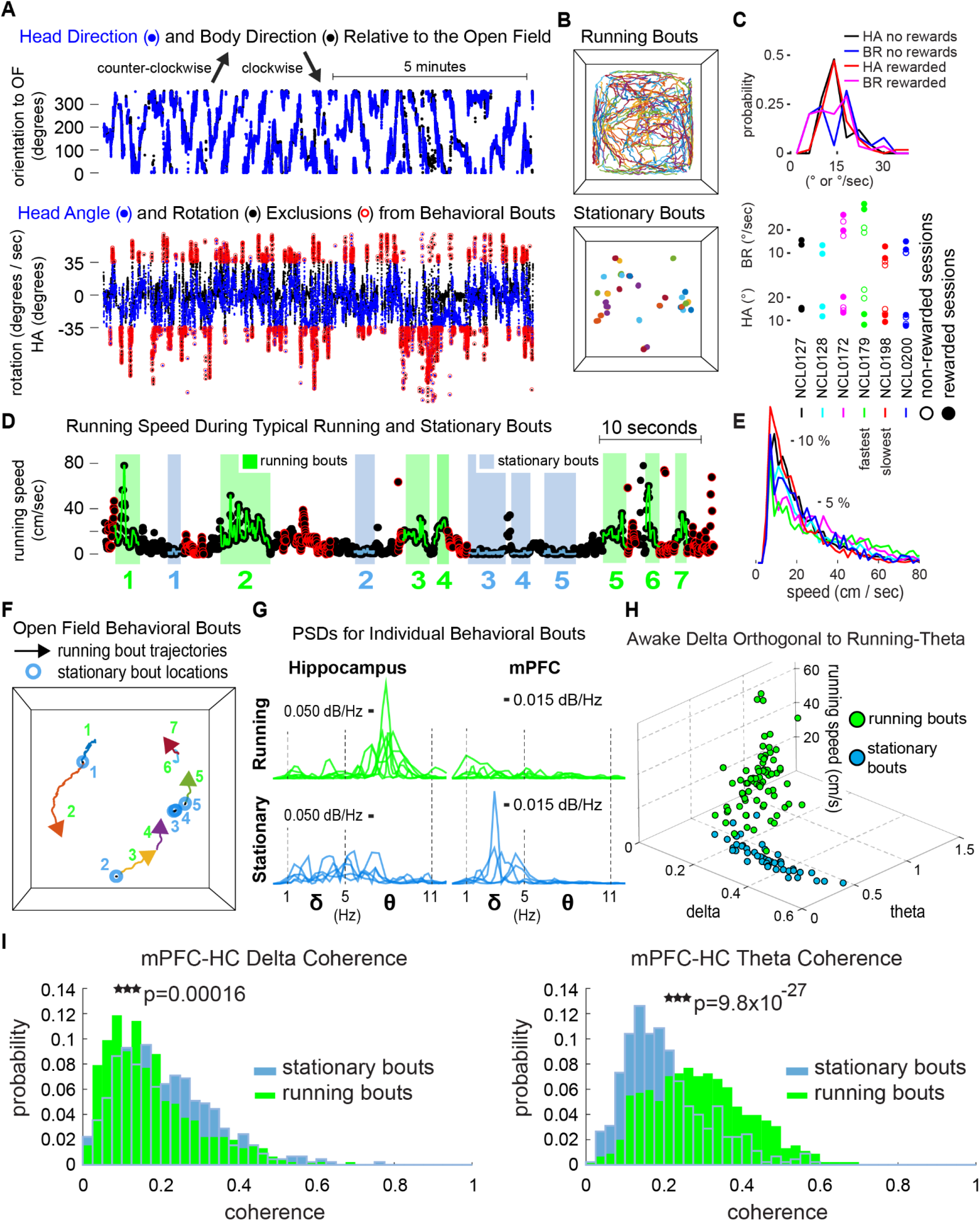
Delta-theta orthogonality and mPFC-HC coherence during intermittent stationary and running bouts. **A.** Representative time-course of body direction (black) and head direction (blue) relative to the open field (**top**). Body rotation speed (BR, black) and head angle (HA, blue) (**bottom**). Data points circled in red indicate time stamps excluded from stationary and running bouts, because they exceeded either 52.5 degrees per second of rotational velocity or 35 degrees of head angle. **B.** Locomotor paths during running bouts (top) and locations of stationary bouts in the open field (bottom) for a representative session. **C**. HA and BR incidence (**top**). Mean HA and BR for each rat (**bottom**). **D**. Intermittent stationary (blue) and running bouts (green), after HA and BR exclusions (red circles) and additional restrictive criteria. **E**. Distributions of speeds visited during running bouts for each rat. **F**. Spatial trajectories of bouts in **D**. **G**. HC and mPFC PSDs for each bout in **D**. HC PSDs during running bouts exhibit prominent theta peaks, whereas delta power was elevated during stationary bouts, particularly in mPFC. **H**. Delta-theta orthogonality divided stationary from running bouts, whereas theta power covaried with speed across running bouts. **I.** Delta coherence between mPFC and HC increased during stationary bouts, and theta coherence increased during running bouts.

## Discussion

We analyzed amplitude and power estimates of delta, theta, and gamma frequency bands of the HC LFP to evaluate network modes and correspondence to behavior in freely navigating rats. As expected, HC theta synchrony covaried with locomotor speed, whereas synchronization in the delta frequency band was minimal during locomotion. Surprisingly, significantly elevated delta power was expressed when animals were stationary even during brief pauses between running bouts.

We used a novel analytic approach to define modes of HC network activity by categorizing power spectra throughout behavior sessions. We present evidence that network modes derived purely from HC activity in this way are strongly related to behavior. Across modes, delta power was orthogonal to theta power paralleling the relationship of each frequency band to locomotor activity. This orthogonality suggests a qualitative difference in network state, rather than an anticorrelation between two continuously varying network properties. We begin to show that spectral modes as we have defined them are functionally distinct by showing differences in intrahippocampal coherence and differences in coherence between mPFC and HC that corresponded to behavioral states. We found that both delta and theta modes were representative of intrahippocampal network coherence independent of clustering strategy and session type. Importantly, our analysis of cross-frequency couplings between frequency bands indicated the possibility that delta-theta, delta-gamma, and theta-gamma relationships may differ both as a function of behavioral state and potentially across recording sites in HC.

Although not addressed by our current analyses, it is a strong hypothesis that there is an anatomical layout in HC wherein delta is stronger in some regions and theta in others, reflecting synaptic input to particular regions within the laminar structure of HC or along the proximal-distal or septo-temporal axes. Such a mapping would be expected to relate to coherence measures between sites within HC and with other structures. To the extent that we evaluated intrahippocampal and mPFC-HC coherence, we found that coherence spectra for pairs of recording sites tended to fall into fairly homogenous groupings of similar coherence spectra that were often markedly distinct from one another (data not shown). We suspect that these groupings delineate important anatomical boundaries, possibly related to limbic thalamic circuitry (Dolleman van der Weel et al., 2019), and thus, application of clustering routines to coherence spectra is likely to be a fruitful approach for further defining network and circuit modes. To this end, it will also be of considerable utility to combine diverse mode definitions, i.e. clustering analysis parameters and clusters derived from different recording sites.

From the present data set it is not possible to determine what cognitive, behavioral, or physiological dimensions might be encoded by the 3-5 fold differences in delta power that was typical, nor by the distribution of power across the delta frequency range (including one or more peaks). On the one hand, one might assume that the degree of synchronization and temporal dynamics of delta-frequency activity are purely epiphenomenal to the integration of up-/down-state transitions from many semi-independent columns across widespread cortical regions that project to the HC. However, it is worth noting how well positioned to influence memory-guided decision making and behavior, or to implement the first steps of episodic memory encoding, are the rapidly-developing delta-dominated network modes amidst an animal’s behavioral and cognitive state trajectories.

Hasselmo and colleagues (2002) suggested that encoding and retrieval mechanisms for episodic memory may be preferentially engaged at different phases of the theta rhythm when different inputs to CA1 are active (Schomburg et al., 2014). Our interpretation of alternating delta and theta modes during behavior is similar, albeit operating at a slower timescale. It could be imagined that these or other mnemonic or cognitive functions are preferentially engaged by alternating locomotor and stationary segments of behavioral sequences and corresponding theta- and delta-dominated circuit dynamics. In such a general formulation, awake delta-dominated states could be taken to reflect some form of meditation on or digestion of recent and ongoing experiences (similar to Todorova and Zugaro, 2019), or decision making and planning for upcoming actions. Behaviorally, locomotor pauses can segment sequences of actions or events within an episodic organization. Could awake delta-dominated modes contribute to segmentation of episodic memory? Is there a predominance of cortical silence during awake delta modes as had been previously been ascribed to delta waves during slow wave sleep? Or do prospective or retrospective representations exist during awake delta modes that could be of particular importance for encoding aspects of ongoing experience to guide future behavior?

## Acknowledgements

This work was supported, in part, by R01 ES006189 to T.R.G., R01 MH113626 to T.A.A. and funding from the Feinberg Foundation provided to T.A.A.

## Supplemental Figures

**Supplemental Figure 1.**
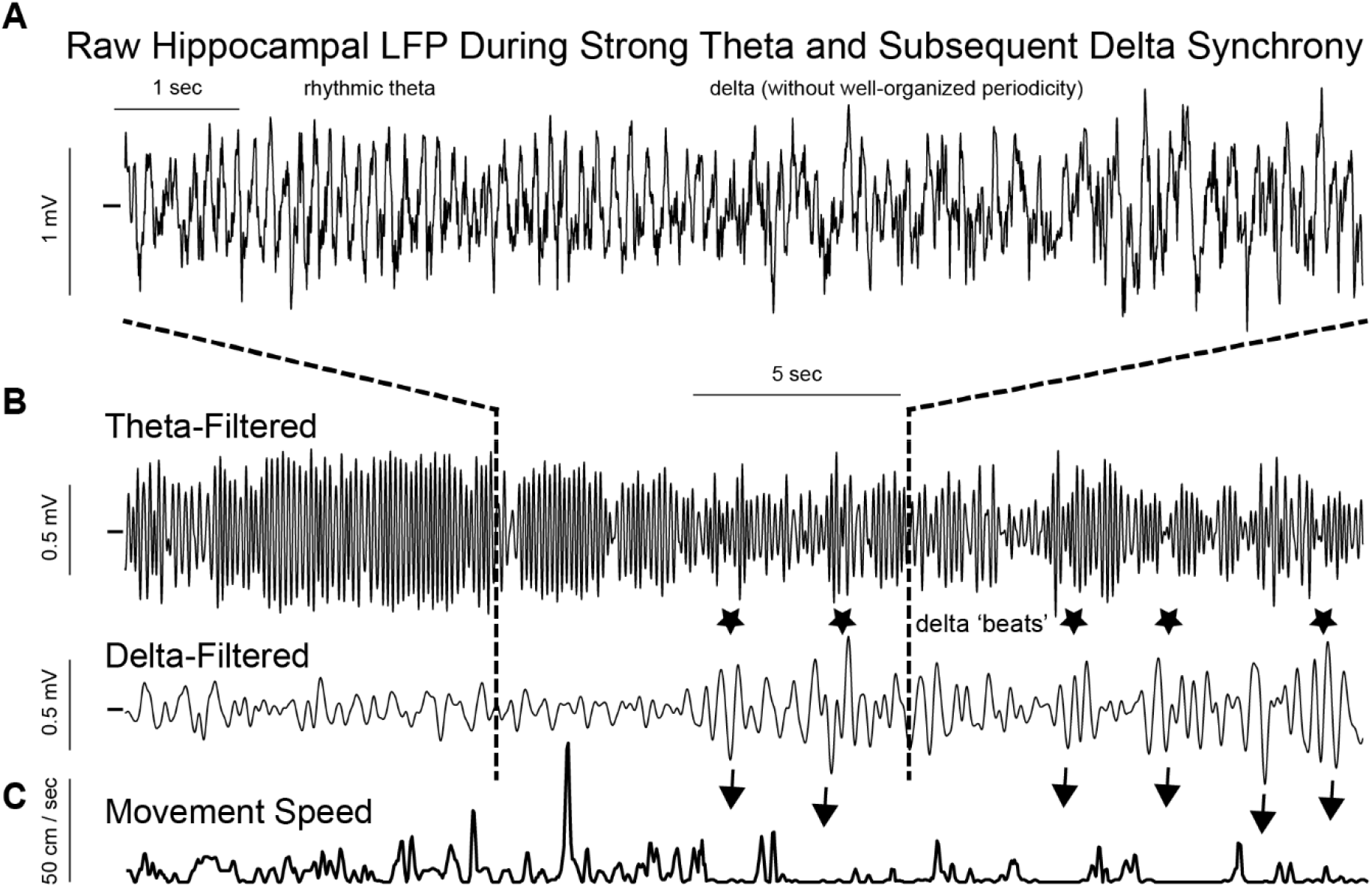
Alternating periods of theta and delta during intermittent running. **A.** HC LFP showing strong theta during the first half and strong delta afterwards. **B.** Theta band-pass and delta band-pass filtered traces of the LFP in **A**. Note the appearance of ‘delta beats’ (stars) with relatively large amplitudes, corresponding to periods when the rat was very still (arrows) and punctuated by brief movements (**C**).

**Supplemental Figure 2.**
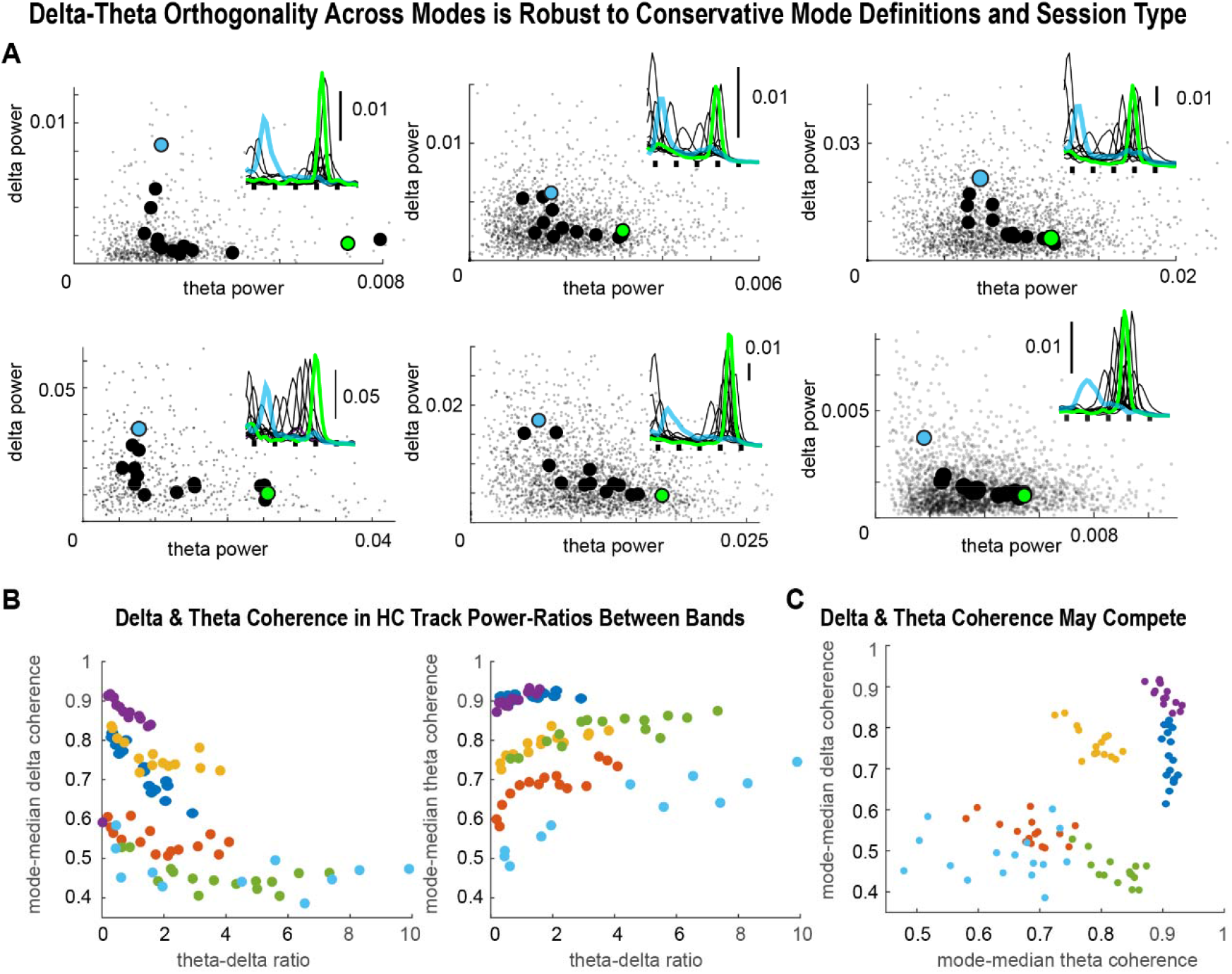
Robustness of delta-theta orthogonality and intrahippocampal coherence. **A.** Delta and theta power were orthogonal across spectral modes derived using normalized windowed PSDs from rewarded sessions for each rat. Small grey points indicate delta and theta power for each windowed PSD. Large black, blue, and green points plot power measures for each mode. Spectral modes, i.e. average PSDs, are inset. Blue and green points in power scatterplots correspond to inset modes of the same colors. **B**. Median delta (**left**) and theta coherence (**right**) among six pairs of simultaneously-recorded HC LFPs averaged across all instances of each mode. Intrahippocampal delta coherence decreased monotonically, while theta coherence increased with increasing theta-delta ratios across modes. **C**. Intrahippocampal delta and theta coherence may be negatively correlated with slopes specific to each rat.

